# Unification of Environmental Metabolomics with Metacommunity Ecology

**DOI:** 10.1101/2020.01.31.929364

**Authors:** Robert E. Danczak, Rosalie K. Chu, Sarah J. Fansler, Amy E. Goldman, Emily B. Graham, Malak M. Tfaily, Jason Toyoda, James C. Stegen

**Affiliations:** Pacific Northwest National Laboratory, Washington, USA; Environmental Molecular Sciences Laboratory, Washington, USA; Department of Soil, Water and Environmental Science, University of Arizona

## Abstract

Environmental metabolomics, enabled by high-resolution mass spectrometric techniques, have demonstrated the biogeochemical importance of the metabolites which comprise natural organic matter (NOM). However, significant gaps exist in our understanding of the spatiotemporal organization of NOM composition. We suggest that the underlying mechanisms governing NOM can be revealed by applying tools and concepts from metacommunity ecology to environmental metabolomics. After illustrating the similarities between metabolomes and ecological communities, we call this conceptual synthesis ‘meta-metabolome ecology’ and demonstrate its potential utility using a freshwater mass spectrometry dataset. Specifically, we developed three relational metabolite dendrograms using combinations of molecular properties (i.e., aromaticity index, double-bond equivalents, etc.) and putative biochemical transformations. Using these dendrograms, which are similar to phylogenetic or functional trait trees in ecological communities, we illustrate potential analytical techniques by investigating relationally-informed α-diversity and β-diversity metrics (e.g., MPD, MNTD, UniFrac), and null model analyses (e.g., NRI, NTI, and βNTI). Furthermore, we demonstrate that this synthesis allows ecological communities (e.g., microbes) and the metabolites they produce and consume using the same framework. We propose that applying this framework to a broad range of ecosystems will reveal generalizable principles that can advance our predictive capabilities regarding NOM dynamics.

## Introduction

Environmental metabolomics enables the investigation of the metabolic processes and interactions occurring within an ecosystem and can provide deep insight into ongoing biogeochemical cycles (Graham *et al*. 2018; Stegen *et al*. 2018; Sengupta *et al*. 2019; Garayburu-Caruso *et al*. 2020). This knowledge has been collected using high-resolution mass spectrometric techniques, such as like Fourier transform ion cyclotron resonance mass spectrometry (FTICR-MS) and Orbitrap, which have allowed researchers to investigate the individual carbon compounds that constitute natural organic matter (NOM). As these studies increasingly become spatiotemporally resolved, an investigation of the underlying processes driving metabolome variability becomes necessary in order to develop transitive principles and enhance predictive capabilities across ecosystems. While many metabolomics studies have used multivariate methods to identify if differences exist between metabolomes (Kellerman *et al*. 2014, 2019; Dalcin Martins *et al*. 2017; Graham *et al*. 2017; Tfaily *et al*. 2018; Zark & Dittmar 2018), they have limited capacity to reveal processes that constrain or promote variation (Gilbert *et al*. 2012; Stegen *et al*. 2012; Cavaco *et al*. 2019). To better understand the processes governing metabolome composition, we propose integrating concepts and tools developed in metacommunity ecology, the study of communities across scales (Leibold *et al*. 2004), with environmental metabolomics. This will allow us to investigate mechanisms underlying spatiotemporal dynamics of metabolites as a conceptual analog to ecological metacommunities, and ultimately uncover transferable principles related to NOM organization. Our specific goals are to (1) explore the conceptual parallels between environmental metabolite assemblages and ecological metacommunities and (2) demonstrate how concepts and analysis tools from metacommunity ecology can be used to study environmental metabolites using an example FTICR-MS dataset from the Columbia River and adjacent riverbed. We demonstrate that novel insights are revealed by applying metacommunity ecology theory to environmental metabolomics and contend that this constitutes the new line of inquiry called ‘meta-metabolome ecology.’

### Conceptual parallels between metabolites and ecological units

Ecological communities are assembled through the collective action of random birth and death events, dispersal of individuals, deterministic factors which affect the relative fitness of taxa, and the evolutionary processes of diversification and extinction (Vellend 2010). Therefore, any community can be assumed to be the outcome of many different ecological and evolutionary assembly processes experienced throughout time and space (Nemergut *et al*. 2013). Similarly, ecosystem metabolomes (i.e., assemblages of ‘metabolites’ which are discrete organic compounds that can be subject to biological activity) represent the collective outcome of historical processes that have resulted in the gain, loss and transformation of individual metabolites. We suggest that this set of historical processes share many parallels with those governing ecological community assembly.

Deterministic factors which influence ecological community composition occur due to systematic differences in birth and death rates among resident taxa (i.e., some taxa more successfully produce offspring than others given some environmental constraint) (Fine & Kembel 2011; Kraft *et al*. 2011; Stegen *et al*. 2012, 2013). For example, differential abilities across taxa to scavenge nutrients can lead to a deterministic change in community structure through time (Cavender-Bares *et al*. 2009). Similarly, individual metabolites within metabolite assemblages will undergo fluctuations in production rate (analogous to birth rate) and degradation rate (analogous to death rate), driven by abiotic or biotic transformations (Cory & Kling 2018; Graham *et al*. 2018; Kellerman *et al*. 2019; Garayburu-Caruso *et al*. 2020). Any factors that adjust these rates could lead to a deterministic shift in the composition of the metabolite assemblage. For example, the preference of microorganisms for organic nitrogen in a nitrogen-limited environment could lead to a deterministic shift in metabolite composition such that biogeochemical hotspots become characterized by nitrogenous metabolites (Graham *et al*. 2018). The persistence of low oxidation state organic carbon within anaerobic environments due to the preferential consumption of more thermodynamically available carbon represents another potentially deterministic shift (Boye *et al*. 2017).

Aside from purely deterministic environmental pressures, dispersal processes can strongly influence ecological community composition through variations in organismal exchange rates. When the rate of exchange between two separated communities is high, these communities can become more homogeneous with respect to each other due to ‘mass effects’ (e.g., the flow of individuals from high to low population size) (Shmida & Wilson 1985; Holyoak *et al*. 2005; Urban *et al*. 2008). Although metabolites are governed by passive movement, high rates of metabolite exchange (e.g., via advective hydrologic transport) could conceivably homogenize assemblages through space. At the other extreme, dispersal rates can be low enough that ecological communities diverge in composition as a result of stochastic ecological drift (i.e., random fluctuations in birth and death rates). If there is a lack of temporally or spatially consistent factors driving variation in metabolite production/degradation rates, a dynamic similar to ecological drift should emerge, referred to here as ‘metabolite drift’ where unstructured compositional deviations occur in metabolite assembles. For example, if dispersal is limiting and any thermodynamic or nutritional requirement is too weak, we could expect a ‘metabolite drift’ signal. Therefore, low exchange rates of metabolites can lead to outcomes that are conceptually analogous to dispersal limitation in ecological communities.

Despite the many parallels between ecological communities and metabolite profiles, there are important differences. For example, no clear parallels can be drawn between the evolutionary processes influencing ecological communities and metabolite assemblages. Specifically, the rates of diversification and extinction which affect the relationship between the regional and local species pools do not translate to metabolites (Vellend 2010). This is because metabolites cannot reproduce or pass along heritable traits. Instead, evolutionary processes could exert an indirect effect on the metabolome by acting upon biological taxa participating in production or degradation of individual metabolites. While we focus on conceptual parallels to deterministic and stochastic ecological factors, we can envision an extension of our framework which integrates indirect evolutionary forces into understanding spatiotemporal dynamics of ecosystem metabolomes.

Species interactions are another feature for which there are discrepancies between ecological communities and metabolite assemblages. In some situations, there is good correspondence. For example, in both ecological communities and metabolite assemblages, members (i.e., biological species or metabolites) will exert indirect and direct pressures on other members, through interactions like predation of individuals or the complexation of metabolites, respectively (Shurin & Allen 2001; Lehmann & Kleber 2015; Xue & Goldenfeld 2017). However, unlike ecological communities, metabolite assemblages are incapable of competitive exclusion such that mechanisms of coexistence don’t have direct analogs within metabolite assemblages (Vellend 2010). Despite these departures from traditional ecological communities, we assert that examining metabolite profiles using metacommunity ecological concepts and tools provides a new perspective on ecosystem metabolites that will enable new conceptual and mechanistic understanding. We further assert that this knowledge is a critical element of ongoing efforts to improve process-based predictive models of ecosystem function (e.g., reactive transport models of river corridors or soil carbon models) that underlie broader Earth system function.

## Extending ecological analyses to metabolite assemblages

Given the similarities between the deterministic and stochastic processes acting upon ecological communities and metabolite assemblages, we propose that ecological concepts and tools can be applied to metabolomes to gain insight into factors governing metabolite spatiotemporal dynamics. We specifically attempt to understand the assembly processes that govern the composition of metabolite assemblages because they are directly analogous to ‘community assembly processes’ that govern the composition of ecological metacommunities. We focus on applying phylogeny-informed diversity metrics and phylogenetic-based null models because traditional multivariate statistics are not sufficient to infer assembly processes (Chase & Myers 2011; Stegen *et al*. 2012, 2013). The null modeling approach adapted here for metabolite assemblages has been shown to provide robust estimates for the relative influences of different assembly processes (Stegen *et al*. 2012, 2013, 2015, 2016a; Graham *et al*. 2016; Danczak *et al*. 2018). Due to the absence of explicit metabolite phylogeny, we use trait-based dendrograms that represent shared and divergent metabolite properties (Swenson *et al*. 2012). As in ecological functional trait or phylogenetic α-diversity and β-diversity analyses, the dendrogram approach provides information beyond simple taxonomic (or metabolite) assignments (Kraft *et al*. 2007; Chase *et al*. 2011; Tucker *et al*. 2016). We demonstrate how relational dendrograms can be used to study metabolome α-diversity and β-diversity, and how they can be used with null model analyses to reveal assembly processes governing the composition of metabolome meta-assemblages.

### An example set of metabolite assemblages and microbial communities

We use metabolite data from the Columbia River corridor to provide an example of how to use a dendrogram-based framework to study the processes influencing metabolite assemblages. In brief, samples of river water and pore water were collected from 5 locations along the Columbia River in Washington State across a ∼1km transect running along the shoreline (see Supplement for details). At each location, filtered river water and subsurface pore water were collected; one replicate of river water was collected, and 3 pore water samples were collected from 30cm depth within a 1m^2^ area using 0.25-inch diameter sampling tubes. Samples were analyzed using FTICR-MS at the Environmental Molecular Sciences Laboratory using previously established methods (see Supplement for details). The raw FTICR-MS data were processed according to established methods to (1) identify peaks from the mass spectra that correspond to unique metabolites identified by their unique mass, (2) calibrate peak/metabolite masses against a standard set of known metabolites, and (3) assign molecular formula based the Compound Identification Algorithm (CIA; see Supplement for details) (Kujawinski & Behn 2006; Tolić *et al*. 2017). Further data analyses are described below in the subsections that use the associated analysis. In addition, water samples were analyzed for basic geochemical parameters (i.e., dissolved organic carbon concentration, specific conductivity, and major anions and cations). We extracted DNA from the filters used to collect aqueous samples and characterized associated microbial communities using 16S rRNA gene sequencing and associated data processing to pick operational taxonomic units and generate a phylogenetic tree (see Supplement for details). All data (i.e., OTU table and FTICR-MS data) can be accessed in the Supplemental Information.

### Building metabolite dendrograms

Tools in metacommunity ecology often leverage relational information such as among-species evolutionary relatedness or functional trait similarities. The associated phylogenetic trees and functional trait dendrograms have enabled researchers to develop metrics which incorporate among-species relational information, leading to novel insights regarding the balance among stochastic and deterministic assembly processes (Faith 1992; Chase *et al*. 2011; Fine & Kembel 2011; Lozupone *et al*. 2011; Stegen *et al*. 2015; Tucker *et al*. 2016). While metabolites do not have genetic sequence information, their characteristics can be approached in a way that is analogous to the functional trait approach in ecological analyses (Swenson *et al*. 2012; Siefert *et al*. 2013). Unlike multivariate dendrograms typically used within metabolomics studies (e.g., Tfaily *et al*. 2018), these dendrograms represent relationships between metabolites and not samples. To this end, we developed and evaluated three methods of measuring trait-like relational information between different chemical compounds using two different information sets: molecular characteristics and biochemical transformations (Figure 1).

**Figure 1:**
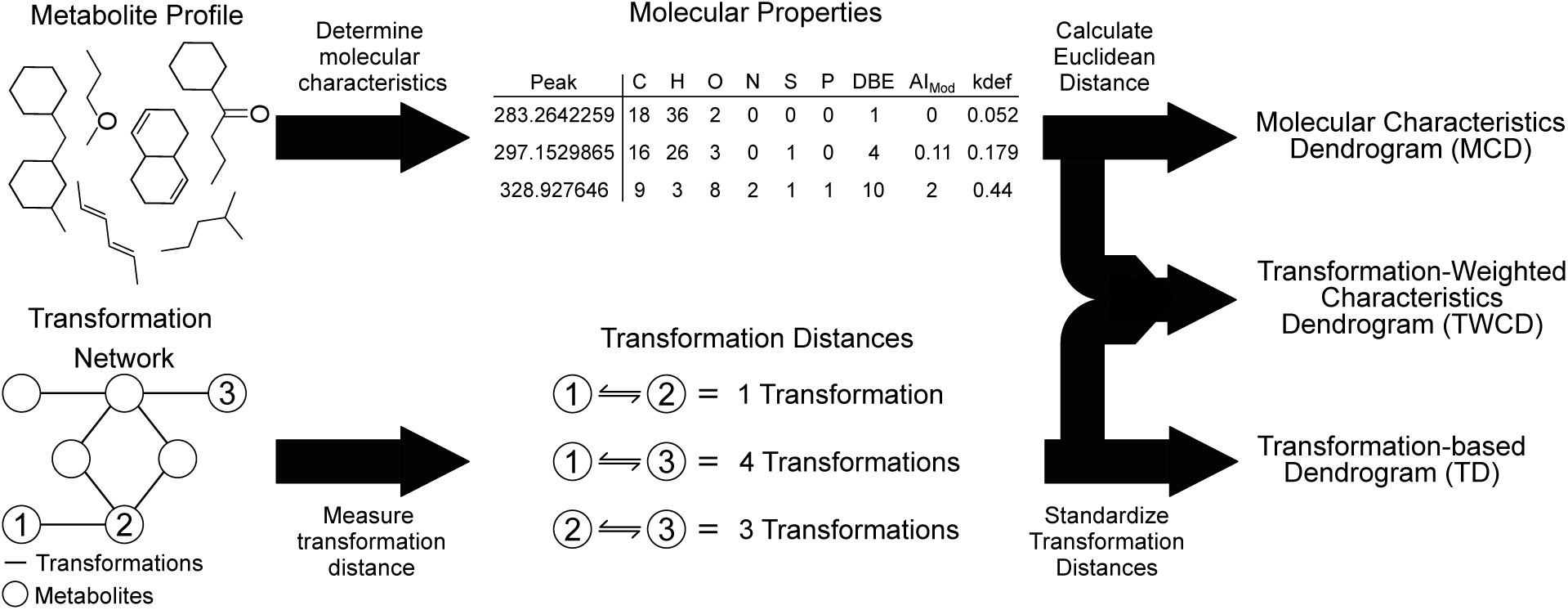
Figure summarizing the steps necessary to create the three dendrograms used throughout this manuscript. The top path (Molecular Characteristics Dendrogram or MCD) demonstrates the relational information provided by molecular properties, like elemental composition and aromaticity index, while the bottom path (Transformation-based Dendrogram or TD) emphasizes the relationships driven by potential biochemical transformation networks. The middle path (Transformation-Weighted Characteristics Dendrogram or TWCD) is a combination of information provided by the top and both paths.

First, we generated a molecular characteristics dendrogram (MCD) which integrates elemental composition (e.g., C-, H-, O-, N-, S-, P-content) and derived statistics (i.e., aromaticity index, double-bond equivalents, etc.) similar to principles outlined in compound classification studies (Hughey *et al*. 2001; Kim *et al*. 2003; Koch & Dittmar 2006; LaRowe & Van Cappellen 2011; Tfaily *et al*. 2015; Bailey *et al*. 2017; Rivas-Ubach *et al*. 2018). Next, we created a transformation-based dendrogram (TD) using putative biochemical transformations identified by aligning mass differences to a database of known transformations (Breitling *et al*. 2006; Bailey *et al*. 2017; Graham *et al*. 2017, 2018; Moritz *et al*. 2017; Stegen *et al*. 2018; Sengupta *et al*. 2019). Finally, we made the transformation-weighted characteristics dendrogram (TWCD) which is a combination of the MCD and TD. Detailed differences between these dendrograms are explored in the Supplement, but each resulted in different metabolite clustering patterns that help provide deeper insight into ecosystem assembly. We suggest that while other approaches to estimating dendrograms from metabolite data exist, the MCD, TD, and TWCD provide a complementary set of methods that are useful for studying the spatiotemporal organization of metabolite assemblages.

### Using metabolite dendrograms to study metabolite diversity and assembly processes

From a practical perspective, the three dendrograms provide a foundation for studying metabolite assemblages with ecological tools that traditionally use phylogenetic or functional trait data. For example, below we show how metabolomes can be studied using metrics associated with richness (Faith’s PD, UniFrac), overall divergence (MPD), and nearest neighbor divergence (MNTD) (Faith 1992; Webb *et al*. 2002; Lozupone *et al*. 2011; Tucker *et al*. 2016). As a parallel to ecological analyses, these metrics can be used to study the spatial and temporal organization of metabolome meta-assemblages.

Many ecological studies track trait dynamics or utilize identity-based (i.e., taxonomic) analyses such as Bray-Curtis dissimilarity to infer ongoing ecosystem processes (Bray & Curtis 1957; Shipley *et al*. 2006). There are, however, exciting opportunities to go further by using additional tools from metacommunity ecology that are designed to infer and quantify assembly processes. Null models represent one set of tools that provide additional insight and complement traditional α-diversity and β-diversity analyses. By applying commonly used phylogenetic null models, we can investigate the processes responsible for structuring metabolite assemblages. First, to assess whether α-diversity was more or less structured that would be expected by random chance, we calculated both the net relatedness index (NRI) and nearest taxon index (NTI), which are null models for MPD and MNTD respectively (Webb *et al*. 2002; Fine & Kembel 2011). For both these metrics, positive values indicate clustering within the dendrogram while negative values signify over-dispersion (Webb *et al*. 2002).

Ranging from cold weather adaptation in forests (Feng *et al*. 2015), labile carbon degradation in bacterial communities (Goldfarb *et al*. 2011), or host range/soil adaptations in root-associated mycobiomes (Schröter *et al*. 2019), these metrics have revealed patterns in phylogenetic trait conservation through different phylogenetic lineages (Martiny *et al*. 2013). Despite examining different ecosystems and scales, a common framework enabled researchers to develop consistent conceptual conclusions. In turn, these null models could provide a similar framework for metabolite assemblages dependent upon the dendrogram. For example, overdispersion observed on the MCD might suggest broadly distributed thermodynamic properties while it could indicate biochemically disconnected peaks on the TD. Such analyses will allow researchers to ask and answer new questions regarding the development of metabolite assemblages.

To further explore the ecological assembly processes structuring metabolite profiles, we calculated the β-nearest taxon index (βNTI; detailed extensively in Stegen *et al*. 2012, 2015). This metric compares the observed β-mean nearest taxon distance (βMNTD) between two communities to a null expectation generated by breaking observed dendrogram associations. While typically informed using abundance data, this null model still produces useful information with presence/absence data. When a comparison between two ecological communities significantly deviates from the null expectation (indicated by |βNTI| > 2), we infer that some deterministic process is responsible for the observed pattern. These deterministic processes can be further separated into those which drive a divergence between communities, termed ‘variable selection’ (indicated by βNTI > 2), and those which drive a convergence between communities, termed ‘homogeneous selection’ (indicated by βNTI < −2). If the pairwise comparison instead mirrors the null expectation (indicated by |βNTI| < 2), we infer that stochastic processes drive observed differences. These stochastic processes can be further distinguished using an identity-based (“taxonomic”) null model, like Raup-Crick, which is able to disentangle dispersal limitation from homogenizing dispersal (i.e., mass effects) when used in conjunction with βNTI.

Previous studies that combined βNTI and Raup-Crick have revealed significant variation in the relative influences of different community assembly processes among systems. This previous work spans a broad range of systems such as the subsurface (Stegen *et al*. 2013, 2015; Graham *et al*. 2016, 2017), soils (Dini-Andreote *et al*. 2015; Bottos *et al*. 2018; Feng *et al*. 2018; Sengupta *et al*. 2019), human microbiome (Martínez *et al*. 2015), and marine (Wu *et al*. 2018). While each study is framed around a distinct set of questions, they are united by an emphasis on understanding the relative contributions of different assembly processes and linking assembly processes to other system features (e.g., redox conditions, succession, abiotic dynamics, ecosystem function, human society, etc.). Having a common conceptual grounding across studies provides opportunities to develop theory that applies across systems and to investigate assembly processes affecting metabolite assemblages. Moreover, this common theory can be used to study ecological communities (e.g., microbes) and the metabolites they transform using the same framework. It will be possible to evaluate the degree of coordination between ecological and metabolite assembly processes and use associated outcomes to inform the mechanistic models that represent organisms and metabolites within dynamic ecosystems (e.g., reactive transport models).

## Example use of the dendrogram-based framework

We next use data from the Columbia River corridor to provide an example of how to use the dendrogram-based framework to study the processes influencing metabolite assemblages. Samples were collected from different environments (i.e., river water and subsurface pore water) but in a system with significant hydrologic connectivity between these environments (Stegen *et al*. 2016b). We use the FTICR-MS data discussed earlier to explore within-assemblage diversity (i.e., α-diversity) and between-assemblage compositional differences (i.e., β-diversity). These analyses are pursued with and without dendrogram-based quantification to compare insights between traditional approaches and dendrogram-enabled analyses. In addition, we primarily use dendrogram-based null modeling (e.g., βNTI) to investigate assembly processes, though we later combine it with dendrogram-free null modeling (e.g., Raup-Crick) to expand our conclusions. We expect that the distinct river water and pore water environments will lead to deterministic signatures associated with variable selection while the hydrologic connectivity will manifest as stochastic signatures associated with homogenizing dispersal due to advective transport and metabolite mixing. In addition, we show how null model outcomes associated with metabolite assemblages can be related to null model outcomes associated with microbial communities. This provides an opportunity to evaluate the degree to which assembly processes are coordinated between microbial communities and the metabolites they produce and consume. This represents a conceptual unification across ecological communities, the resources they depend on, and the influences they have over environmental systems.

It is important to recognize that the sample set used here is for demonstration purposes and is therefore relatively small to facilitate straightforward interpretation. We expect significantly different analysis outcomes when the methods developed here are applied to other sample sets and other environmental systems. For example, while we do not observe clear associations between microbial community and metabolome assembly processes, such associations may be stronger in other systems and/or in larger sample sets spanning a broader range of conditions.

### Deterministic organization of metabolome α-diversity is revealed using among-metabolite relational information

Many metabolomic studies employ common multivariate statistics (e.g., ordinations) to determine whether differences exist between samples or sample groups (Dalcin Martins *et al*. 2017; Tfaily *et al*. 2018; Zark & Dittmar 2018). While this can provide useful insights into similarities between samples, these methods do not incorporate among-metabolite relational information. Just as in ecological metacommunities, integrating relational information (e.g., phylogenetic or functional trait relationships) expands the breadth of inquiries one can pursue. The dendrogram-based approach developed here allows relationally-informed α-diversity metrics to be applied to metabolite assemblages and can be used to investigate patterns driven by shared molecular characteristics and biochemical transformations.

Dendrogram Diversity (DD), directly analogous to Faith’s Phylogenetic Diversity (Faith 1992), is a relationally informed metric that quantifies the total dendrogram branch length occupied by a given metabolome. Higher values indicate metabolomes that span a broader range of molecular properties (MCD), that span more broadly across the biochemical transformation network (TD), or both (TWCD). The TWCD values for DD were significantly higher for pore water than river water, but the MCD- and TD-based DD did not differ between pore and river water (Figure 2b). Two additional α-diversity metrics, mean pairwise distance (MPD) and mean nearest taxonomic distance (MNTD), revealed that pore water and river water metabolites share similar dendrogram topologies (i.e., compounds within a given group have similar branch lengths to other compounds) (Figure 2cd).

**Figure 2:**
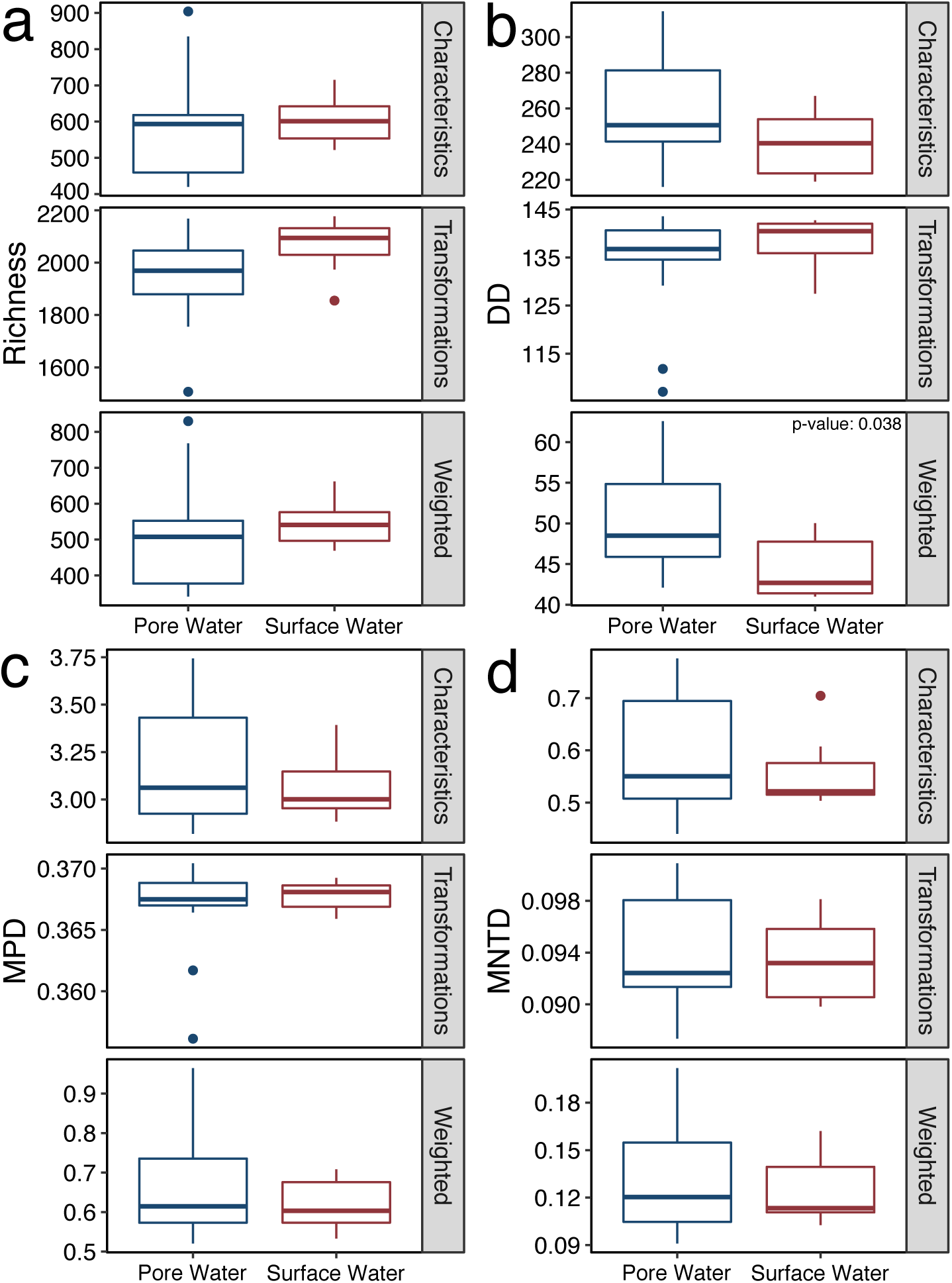
Alpha diversity results for the metabolite data. *a*) Richness (akin to metabolite count). *b*) Dendrogram Diversity (DD) which is analogous to Faith’s Phylogenetic Diversity (PD). *c*) Mean Pairwise Distance (MPD). *d*) Mean Nearest Taxon Distance (MNTD). If differences between surface water and pore water samples was significant as determined by a Mann Whitney U test, the p-value is indicated within the plot.

These dendrogram-based α-diversity metrics indicate that pore water metabolites are slightly more diverse in that they span a broader range of molecular properties. However, pore and river water metabolites are equally diverse with respect to potential biochemical transformation connections. This highlights that multiple dimensions of diversity exist within ecosystem metabolomes, each providing a different window into metabolome organization. The combination of dendrogram-free (e.g., number of unique metabolites) and dendrogram-based (e.g., DD) analyses provides an approach to investigate more dimensions of metabolome diversity than would be possible otherwise. Rather than focusing purely on between-group differences in molecular stoichiometry or other properties (e.g., thermodynamics), relationally informed α-diversity metrics allow new questions related to the organization of metabolite assemblages to be assessed. For example, “How consistent are metabolite richness and DD?” or “How do DD values obtained from MCDs compare to those from TDs across systems?” can now be interrogated. General patterns which emerge from cross-system analyses could point to transferable principles that could be new structural elements integrated into mechanistic models linking metabolite chemistry to microbial and biogeochemical function.

To go beyond direct characterization of α-diversity, null modeling is often used in community ecology to evaluate whether community structure deviates from a stochastic expectation (Gotelli & Colwell 2001). A broad range of informative null model analyses can be used when a phylogenetic or relational dendrogram is available. In addition, α-diversity phylogenetic null models provide opportunities to evaluate questions not accessible via analyses that do not use relational information. Outcomes can be used in a variety of ways, such as revealing whether a given community is phylogenetically over- or under-dispersed (Webb *et al*. 2002; Fine & Kembel 2011; Tucker *et al*. 2016). Common null models like net relatedness index (NRI) or nearest taxon index (NTI) can be extended to metabolite assemblages through the use of one or more metabolite dendrograms. Analogous to applying these methods to ecological communities, the degree of clustering or over-dispersion in metabolite assemblages can be quantified.

Both NRI and NTI revealed that the pore water and river water metabolite assemblages had significantly more clustering than would be expected by random chance as indicated by high positive values. This was consistent across all three dendrograms and indicates an important influence of deterministic assembly processes that constrain the composition of metabolite assemblages. Furthermore, river water metabolites had greater clustering than pore water in every analysis aside from the MCD- and TD-based NTI (Figure 3). Interpreted in concert with the α-diversity patterns discussed above, the null model results suggest that the decreased TWCD-based DD within river water metabolites is driven by an increased amount of tip-level clustering. Greater tip-level clustering in the TWCD indicates the presence of metabolite groups that are highly similar to each other in terms of their molecular properties and shared biochemical transformations.

**Figure 3:**
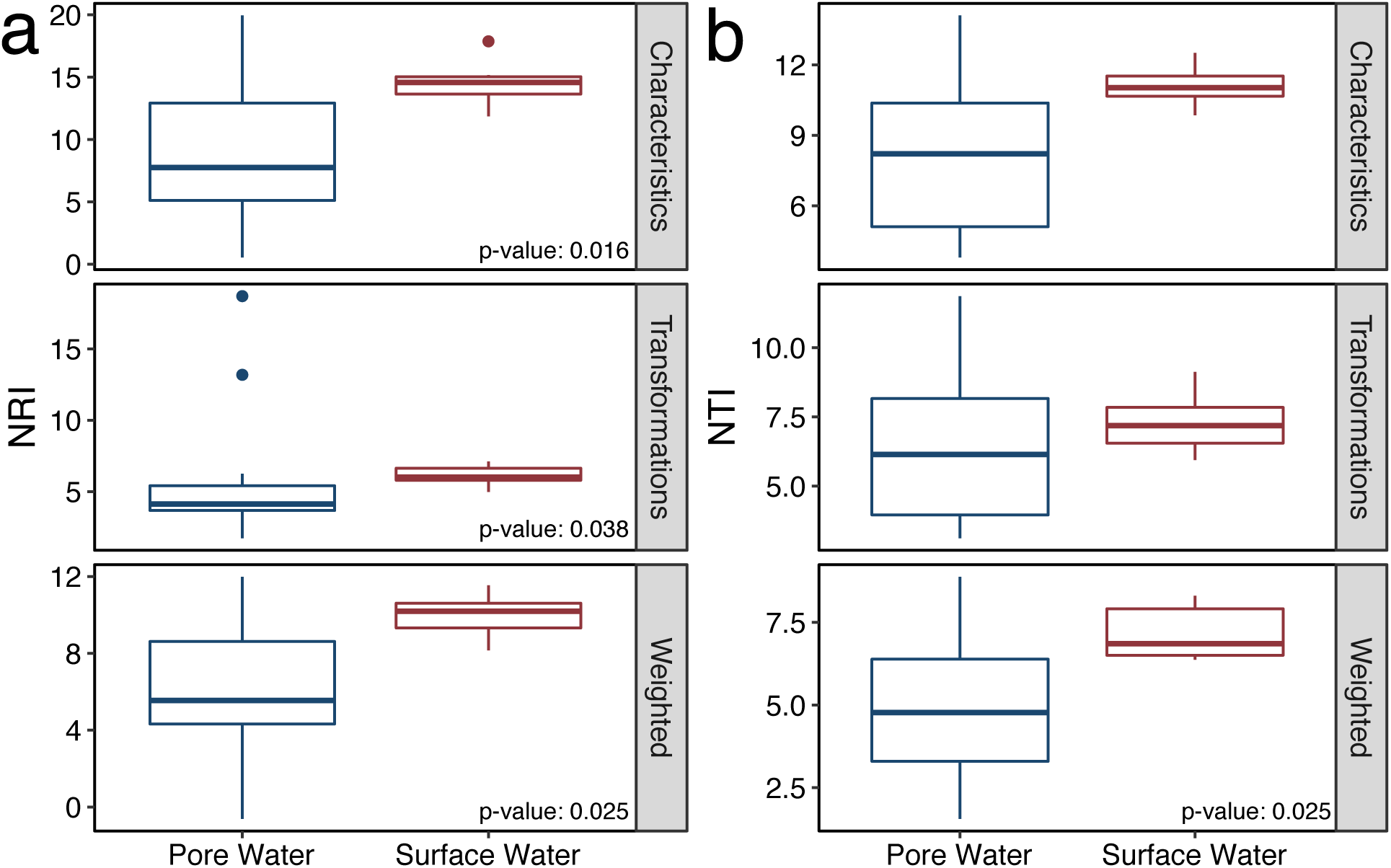
Alpha diversity null modeling results for the metabolite data. *a*) Net Relatedness Index (NRI). *b*) Nearest Taxon Index (NTI). If differences between surface water and pore water samples was significant as determined by a Mann Whitney U test, the p-value is indicated within the plot.

Examining metabolite assemblages through the lens of community ecology provides opportunities to generate conceptual outcomes that would not be possible with traditional multivariate analyses. Here, combining outcomes across α-diversity analyses revealed both pore and surface water were deterministically organized despite divergent mechanisms. More specifically, pore water metabolites were more over-dispersed than the river water metabolites according to every NRI and one NTI analysis. This demonstrates that pore water metabolites (1) span a broader range of molecular properties and (2) are separated by a larger number of biochemical transformations. Systematic differences in the molecular properties and biochemical transformation networks topologies between pore and river water indicates that different localized processes influence metabolite assemblages across the river corridor ecosystem. This suggests a need to understand variation in the mechanisms that underlie metabolome assembly processes. To this end, additional insights can be gained by taking further advantage of the conceptual unification of metabolomics and metacommunity ecology to evaluate the processes influencing variation in metabolome composition (i.e., β-diversity).

### Spatiotemporal variation in molecular properties is stochastic, while biochemical relationships are deterministic

As a complement to α-diversity, β-diversity is commonly evaluated to capture multivariate differences among ecological communities. Previous studies have also used dendrogram-free β-diversity metrics (e.g., Jaccard) to study differences among metabolite assemblages (Graham *et al*. 2017; Kellerman *et al*. 2019; Garayburu-Caruso *et al*. 2020). As with α-diversity, these metrics can be extended by utilizing relational information provided by a phylogeny or dendrogram (Fine & Kembel 2011; Lozupone *et al*. 2011; Stegen *et al*. 2013; Tucker *et al*. 2016). This additional relational information enables quantitative evaluation of relative influences of stochastic and deterministic assembly processes influencing spatiotemporal variation in the composition of ecological communities or metabolite assemblages.

While quantitative evaluations of stochasticity and determinism are common within community ecology, they have not been pursued within metabolomics. Such analyses open new conceptual domains focused on the processes causing spatiotemporal variation in metabolomes. For example, while it is well known that stochastic processes work in concert with deterministic processes to influence spatiotemporal variation in ecological communities (Zhou & Ning 2017), it is unknown whether stochastic processes have any significant influence over spatiotemporal variation in metabolite assemblages. Given the strong influence of metabolite assemblages over biogeochemical function (Graham *et al*. 2017, 2018; Stegen *et al*. 2018), any significant influences of stochasticity are likely to alter biogeochemical function in potentially unpredictable ways (Graham & Stegen 2017). Furthermore, given that ecosystem metabolites are both resources for and products of microbial metabolism, strong influences of stochasticity over metabolomes may cascade into microbial community assembly or indicate highly variable microbial metabolic processes at spatial scales below the sample volume. These examples only represent some types of scientific inquiry that can be opened up by examining spatial and/or temporal variation in metabolome composition (i.e., β-diversity) through the lens of meta-community ecology.

To study metabolome β-diversity, we examined both dendrogram-free and dendrogram-based metrics to provide the deepest conceptual insights. Using a dendrogram-free approach, β-diversity results from our dataset revealed greater differences than the *α-*diversity analyses. A Jaccard-based non-metric multidimensional scaling (NMDS) analysis revealed that pore and river water metabolites diverged significantly from each other (Pseudo-F: 1.48, p-value: 0.018; Figure 4a). Incorporating relational information provided by the metabolite dendrograms resulted in the emergence of more defined clusters while maintaining significant differences. Here, relational information was integrated using unweighted UniFrac, which compares the number of shared and unshared branch lengths between two assemblages (Lozupone *et al*. 2011). Three discrete clusters which were not observed in Jaccard-based analysis emerged when using either the MCD or TWCD, but not the TD (Figure 4). This highlights the deeper level of information provided when considering the molecular and biochemical relationships among metabolites, similar to the use of phylogenetic or functional trait information, and indicates that molecular properties are conserved within particular sets of metabolite assemblages. We infer that there are consistent biotic and/or abiotic processes acting to constrain molecular properties across subsets of metabolite assemblages, but not the biochemical transformations. UniFrac analyses, however, are not capable of identifying the relative contributions of stochastic and deterministic processes which can instead by parsed out through the use of null models.

**Figure 4:**
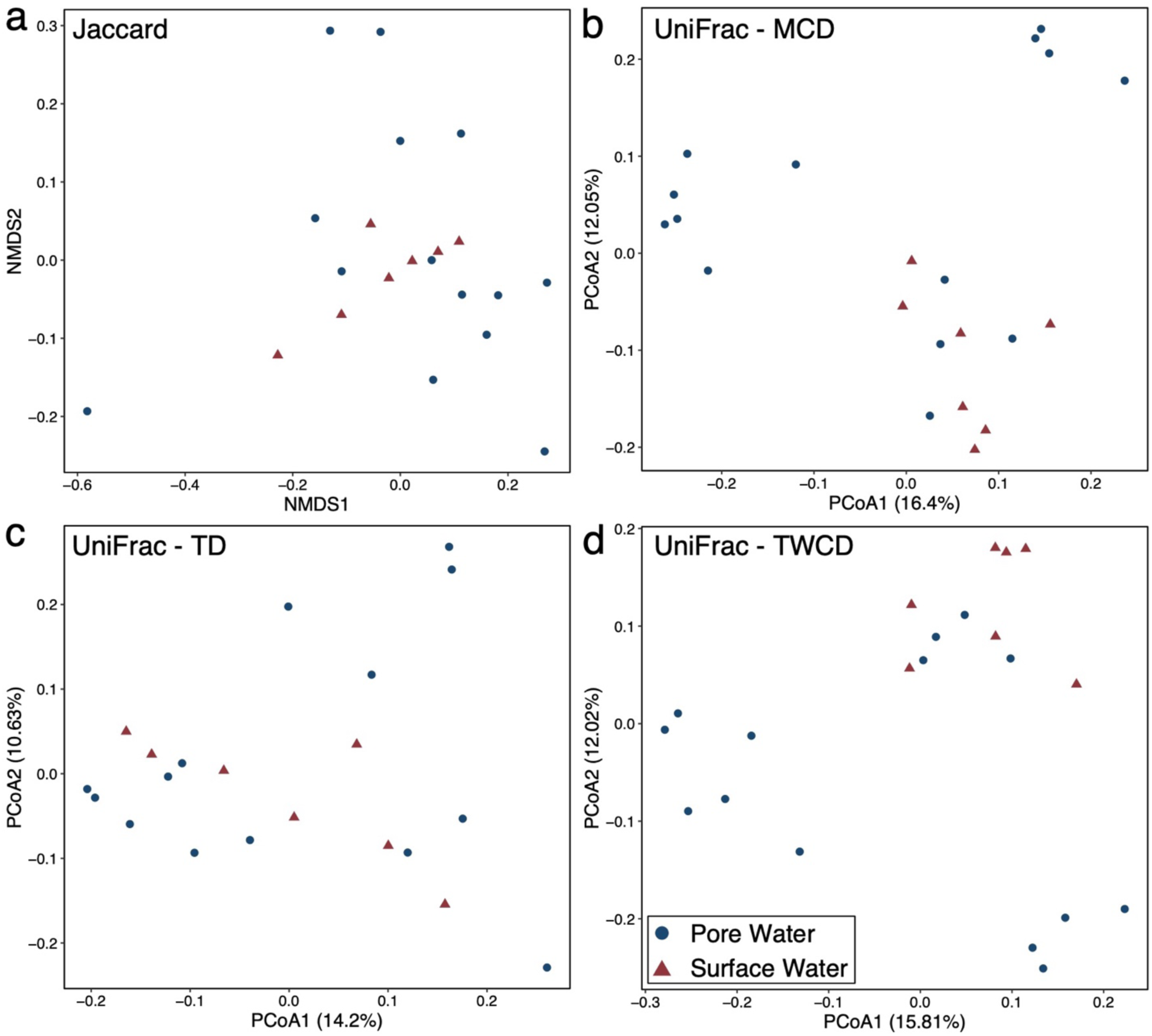
Beta diversity ordinations. *a*) Jaccard Dissimiarlity-based NMDS plot. *b*) UniFrac PCoA generated using the MCD. *c*) UniFrac PCoA generated using the TD. *d*) UniFrac PCoA generated using the TWCD.

Similar to α-diversity, using relational information (e.g., phylogenies or dendrograms) with β-diversity null modeling can reveal the relative influences of stochastic and deterministic processes over spatiotemporal variation in the composition of ecological communities and metabolite assemblages. Given that phylogenetic analyses of microbial communities often use the beta nearest taxon index (βNTI) null model (Danczak *et al*. 2018; Stegen *et al*. 2018; Sengupta *et al*. 2019), we used it here to study metabolite assemblages. We encourage follow-on studies to explore the relative merits of different null modeling approaches applied to metabolome assemblages, however. As reviewed above, βNTI is a pairwise metric and is estimated by quantifying the difference between the observed beta mean nearest taxon distance (βMNTD) and βMNTD expected under stochastic assembly. If a given pair of metabolite assemblages are significantly more different from each other than would be expected under stochastic assembly (indicated by βNTI > 2), we infer that deterministic processes have caused divergence in metabolome composition. This situation is referred to as ‘variable selection’ in the ecological literature. In contrast, if a pair of metabolite assemblages are more similar to each other than expected (indicated by βNTI < −2), we infer that that deterministic processes have constrained the composition of those assemblages to be similar. This situation is referred to as ‘homogenous selection’ in the ecological literature. Lastly, if differences between a pair of metabolite assemblages do not deviate significantly from the stochastic expectation (|βNTI| < 2), we infer that stochastic processes (i.e., mixing, unstructured/inconsistent gains/losses of metabolites, etc.) are primarily responsible for the observed differences in metabolite assemblages. We note that other β-diversity null models could be used to study metabolite assemblages in conjunction with βNTI to reveal additional insights.

Applying the βNTI null model to our dataset revealed that a mixture of homogenous selection, stochastic processes, and variable selection structured spatiotemporal variation in metabolite assemblages (Figure 5a). The influences of these assembly processes differed sharply between molecular property and biochemical transformation-based relationships. βNTI associated with the MCD or TWCD demonstrated that all three structuring processes influenced metabolite assemblages, though stochastic processes and variable selection dominated (Figure 5a). Comparatively, most TD-based βNTI values were > 2, indicating that variation in the topologies of metabolite biochemical transformation networks were predominantly governed by variable selection. As such, the molecular properties in both the pore and river water samples were governed primarily by stochastic processes while the organization of biochemical transformations were deterministic (Figure 5a).

**Figure 5:**
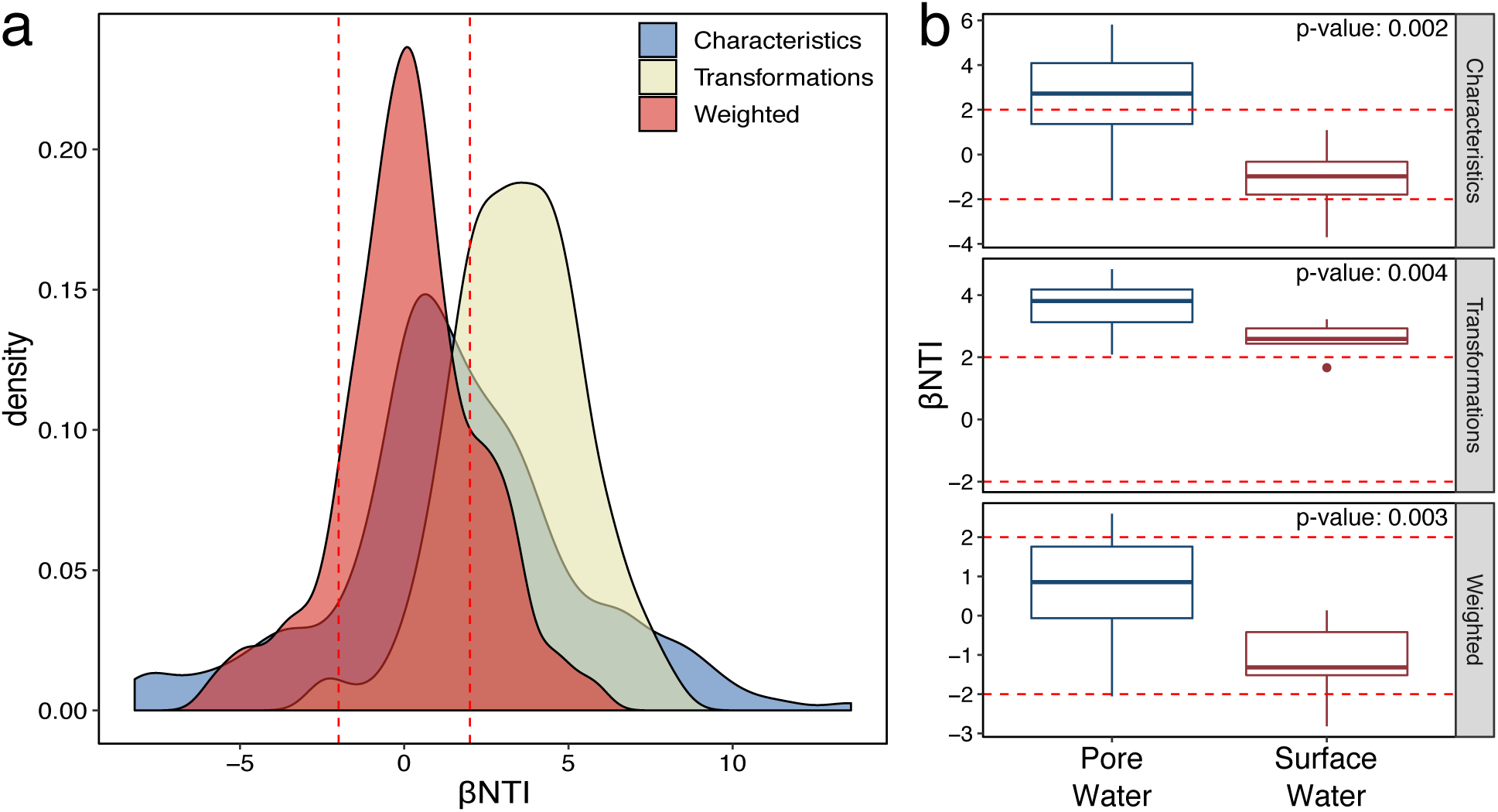
Beta diversity null modeling results. *a*) Density plot of βNTI results for all comparisons. *b*) Boxplots of within-scale βNTI results (e.g., only pore water-to-pore water comparisons). Significant differences as determined by a Mann Whitney U test are indicated by the presence of a p-value within the plot.

### Changing scales leads to additional insights accessible only through null modeling based on relational information

A powerful aspect of null modeling is that one can evaluate the relative influences of stochastic and deterministic influences at different scales (Leibold *et al*. 2004; Swenson *et al*. 2006; Cavender-Bares *et al*. 2009). For example, one can reduce the spatial scale of analysis to study processes causing variation in composition within a given environmental condition and compare that to assembly processes operating at larger scales (i.e., across environments). Here we take advantage of this scale-dependence to study how inferred assembly processes change when constraining analyses within pore or surface water. This provides important insights as there can be processes that drive variation within a given part of an ecosystem that are not responsible for variation across ecosystem components (Cavender-Bares *et al*. 2009). To the best of our knowledge, ecological null modeling is the only robust approach to reveal scale dependence in assembly processes (Stegen *et al*. 2013, 2015).

Our null modeling analyses within pore or surface water indicate that different assembly processes operate within each compartment of the river corridor ecosystem to influence metabolite properties, but not biochemical relationships (Figure 5b). Within pore water, the molecular properties of metabolite assemblages were predominantly governed by variable selection and had higher average βNTI values than the surface water (p-value: 0.002-0.004). This suggests a divergence in localized, deterministic processes across sampled pore water locations. For example, previous work in the study system showed that dynamic (and spatially variable) groundwater-surface water mixing primed microbial respiration (Stegen *et al*. 2018). This is one mechanism among many that could lead to deterministically organized spatiotemporal variation across pore water metabolite assemblages. In contrast, within surface water, the molecular properties of metabolite assemblages were predominantly influenced by stochastic processes. This indicates a potentially strong influence of spatial processes within the surface water, such as significant mixing of metabolite assemblages across the sampled locations.

While assembly processes influencing molecular properties were scale dependent, variable selection consistently governed variation in biochemical relationships. This indicates that each metabolite assemblage in pore or surface water had a distinct transformation network topology. We infer that even when the molecular properties of metabolite assemblages are influenced by stochastic processes (e.g., mixing), the associated biochemical transformations are distinct due to localized processing (e.g., enzymatic degradation) and generation (e.g., metabolite excretion) of organic molecules. By combining the results across scales, we revealed that significant variation within pore water metabolite assemblage assembly processes was not substantial enough to cause deterministic divergence in the molecular properties of surface and pore water metabolite assemblages when analyzed together.

Scale dependence in assembly processes points to a key challenge for representing the properties of metabolite assemblages within predictive models (e.g., integrated hydro-biogeochemical reactive transport models) (Burd *et al*. 2016; Li *et al*. 2017). Such models will need to grapple with when, where, and how to represent detailed mechanisms governing spatiotemporal variation in metabolite assemblages. For example, while the molecular properties of pore water metabolite assemblages diverged from each other due to variation in deterministic processes, it is not clear that variation in those processes is strong enough to warrant representation in predictive models. In contrast, the localized processes appear to strongly influence biochemical transformations at all scales, pointing to the need for representation of associated mechanisms.

As pointed to above, the inferences revealed here through at-scale and between-scale null modeling represent only a few of the conceptual insights that can be gleaned by studying metabolite assemblages through the lens of meta-community ecology. For example, one can parse metabolite assemblages into different kinds of groups based upon elemental composition (e.g., looking at only N containing compounds), thermodynamic properties, or activity within a biochemical network and evaluate variation in assembly processes across these groups. As a demonstration of how insights can be gained by studying assembly processes across metabolite groups, we next examine stochastic and deterministic assembly processes within putatively more or less biochemically active metabolite groups.

### Metabolites across the activity continuum are essential to the deterministic organization of biochemical transformation networks

Metabolites were parsed into putatively more or less biochemically active groups to evaluate whether these groups experience differential assembly processes. This was accomplished by separating those metabolites which were involved in no transformations (less active group) from those which were involved in a comparatively large number (>40) of putative biochemical transformations (more active group). While these groupings are not definitive in terms of relative activity, they provide opportunity for demonstration and preliminary investigation into whether assembly process vary significantly across metabolites that are more or less biochemically connected. One may expect that more active metabolites are more deterministically organized than less active metabolites due to the greater potential for influences of spatial processes over less active members.

Null modeling results from these two groups revealed that putatively more active metabolites experienced stronger deterministic influences. More specifically, the βNTI results from the MCD and TCWD demonstrate that the more active metabolites were influenced by variable selection while the inactive metabolites were stochastically organized (Figure 6). This suggests distinct underlying mechanisms governing the composition of more vs. less active metabolites. The TD-based null modeling, however, showed strong influences of stochastic processes over both more and less active metabolites. This is in direct contrast to the complete metabolite profiles where variable selection drove among-assemblage differences in biochemical relationships (Figure 5). Given that the TD is based upon all possible transformations within a given dataset rather than only those observed in any one sample, the contrasting outcomes suggest that metabolites with relatively few transformation-based connections to other metabolites are key to localized, deterministic organization of biochemical transformations. As such, metabolites across the whole continuum of activity (and connectivity) levels are likely critical to the overall biogeochemical function of the system. Beyond comparing assembly across scales, this framework provides an opportunity to study microbial communities and the ecosystem metabolites they interact with using the same conceptual foundation. For example, one can evaluate the degree to which there is coordination in the assembly processes influencing microbial communities and associated metabolite assemblages.

**Figure 6:**
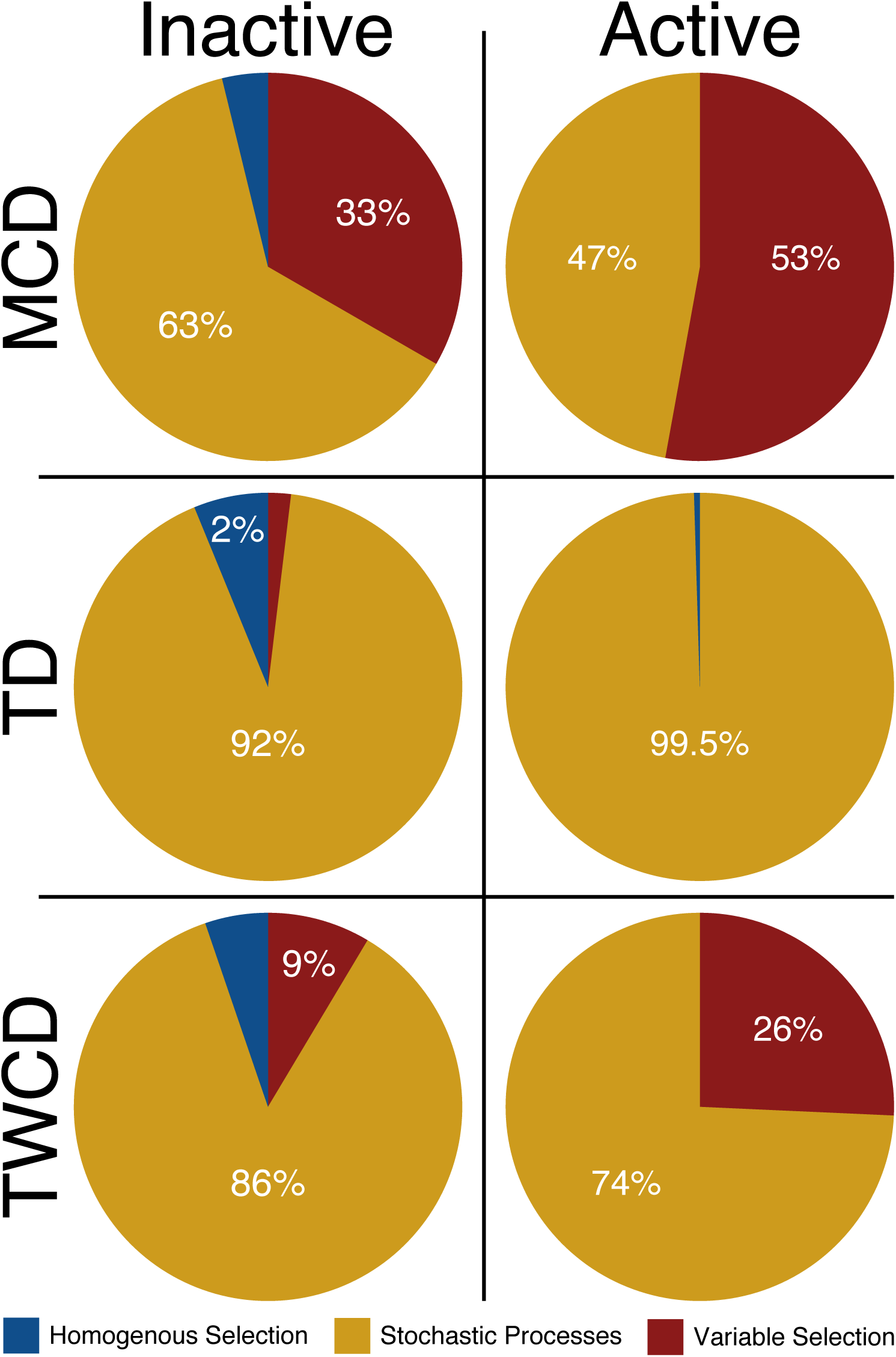
Pie charts illustrating the differences in ecological assembly processes for the active (metabolites involved in >40 transformations) and inactive (metabolites involved in 0 transformations) fractions of the metabolite assemblages.

### Assembly processes influencing microbial communities and metabolites are not coordinated

Microbial communities are a primary driver of ecosystem metabolite transformations and significantly impact rates of organic matter production and degradation (Cory & Kling 2018; Graham *et al*. 2018; Swenson *et al*. 2018; Wieder *et al*. 2018). In turn, there are likely to be dynamic feedbacks between metabolite assemblages and microbial communities (Graham *et al*. 2017, 2018). In order to approximate the extent of these interactions, we examined the relationship between the assembly processes acting upon microbial communities and metabolite assemblages. Given the potential for feedbacks, we expect that the relative influences of assembly processes governing microbial communities will correlate with the assembly processes influencing metabolite assemblages. For example, if the microbial community is shifted due a deterministic process, we expect associated metabolite assemblages will also shift deterministically.

Examining the microbial data in isolation reveals that river corridor communities were equally influenced by variable selection and stochastic processes (∼43% of among-community variation in composition was due to each process). These relative influences mirror those estimated for the metabolite assemblages when using the MCD (∼41% variable selection; ∼46% stochastic). However, once the two models are directly compared to each other using either a Pearson-based or Spearman-based Mantel correlation their correspondence diminished significantly. Specifically, the Pearson-based analysis indicated a weak relationship (Mantel r: 0.298, p-value: 0.067) and the Spearman-based statistic suggested there was no relationship (Mantel r: 0.223, p-value: 0.135).

While the lack of association of assembly processes between microbes and metabolites is contrary to our hypothesis, it points to complex factors influencing both metabolites and microbes. For example, while microbes are one part of the river corridor system that can influence metabolite composition, there are numerous other factors (e.g., vegetation, mineralogy, subsurface hydrology, photooxidation) that likely impact metabolite assemblages (Cory & Kling 2018; Gargallo-Garriga *et al*. 2018; Trusiak *et al*. 2018; Wieder *et al*. 2018; Dwivedi *et al*. 2019). These non-microbial processes likely alter metabolite assemblages in a way that is not reflected in microbial community composition (Gargallo-Garriga *et al*. 2018; Trusiak *et al*. 2018). In addition, metabolite assemblages may change faster than microbial community composition, whereby there may be a closer association between metabolite assemblages and expressed metabolisms (e.g., metatranscriptomes) and/or relative changes in activity (e.g., relative rRNA content across taxa).

The degree to which there are associations between metabolite assemblages and various features of microbial communities is, however, likely to vary through space and time. We posit that new insights can be gained by studying variation in the degree to which assembly processes are coordinated between metabolite assemblages and microbial communities. For example, a strong association between assembly processes influencing metabolites and community composition could indicate a relatively stable system with relatively similar time scales of change for metabolites and microbial composition. Alternatively, strong associations between assembly processes influencing metabolites and expressed microbial metabolism, but not microbial composition, could indicate fast metabolite dynamics coordinated with rapid changes in microbial metabolism despite relatively slow changes in microbial composition. It will be fascinating to explore the associations among assembly processes influencing metabolites and microbial communities and how those associations vary across environmental systems in future studies.

## Conclusions and future directions

Here we have presented a new conceptual paradigm in which metabolite assemblages are treated as analogs to ecological communities and consider this to formalize the sub-discipline of ‘meta-metabolome ecology.’ While the analogy is not complete (e.g., metabolites do not evolve), there are numerous conceptual parallels that allow one to ask new types of questions about environmental metabolites by applying ecology-inspired analyses to metabolite assemblages. We contend that the outcomes of these analyses provide deeper understanding of biochemical and biogeochemical dynamics. For example, identifying when, where, and why metabolite assemblages are governed by deterministic or stochastic processes offers new information about key drivers that may be incorporated into predictive biogeochemical models. This presents exciting opportunities given the integral role of metabolites in emergent biogeochemical function (Graham *et al*. 2017, 2018; Stegen *et al*. 2018). Throughout the analyses discussed above, we have highlighted numerous insights that would not have been possible through traditional multivariate analyses. We anticipate further development of the conceptual, theoretical, and methodological unification initiated here between meta-community ecology and metabolite assemblages.

While we focused heavily on dendrogram-based null models, there are other immediate opportunities to advance understanding by combining dendrogram-based and dendrogram-free null models. This approach has been pioneered in microbial ecology to parse out the relative influences of variable selection, homogeneous selection, dispersal limitation (combined with drift), and homogenizing dispersal (i.e., mass effects). More specifically, this is pursued by combining the βNTI null model with the identity-based Raup-Crick null model. Here, using this approach to refine our understanding of assembly processes influencing metabolite assemblages based upon the MCD revealed that variable selection, homogeneous selection, dispersal limitation (combined with drift), and homogenizing dispersal were responsible for 41.4%, 12.9%, 1%, and 39.5% of variation in pore and surface water metabolite assemblages, respectively. This demonstrates that just as in ecological systems, spatial processes can have significant influences over metabolite assemblages. To fully incorporate metabolites into predictive hydro-biogeochemical models, both selection-based and spatial processes will need to be considered.

Environmental metabolomics has significantly improved our understanding of ecosystem function and biogeochemical cycles (Graham *et al*. 2017, 2018; Stegen *et al*. 2018). By studying metabolite assemblages as analogs to ecological meta-communities, we aim to deepen understanding of the factors influencing the spatial and temporal organization of metabolites within environmental systems. While our demonstration dataset was derived from a river corridor system, the conceptual and methodological framework used here can be applied to any system (e.g., soils, marine, human gut, etc.). We anticipate further development of concepts, theory, and methods that deepen the analogy between meta-communities and metabolite assemblages, but that also diverge from this analogy when needed (e.g., metabolites are not competitively excluded). Application of the framework initiated here to a broad range of ecosystems will provide opportunities to elucidate key principles that are generalizable and that be used to advance our capacity to predict system response to disturbance.

## Acknowledgements

Pacific Northwest National Laboratory is operated by Battelle Memorial Institute for the U.S. Department of Energy under Contract No. DE-AC05-76RL01830. This research was supported by the U.S. Department of Energy (DOE), Office of Biological and Environmental Research (BER), as part of BER’s Subsurface Biogeochemistry Research Program (SBR). This contribution originates from the SBR Scientific Focus Area (SFA) at the Pacific Northwest National Laboratory (PNNL).

## Supplemental Legend

**Supplemental File 1:** Molecular characteristics dendrogram (MCD) generated using the UPGMA hierarchical clustering method.

**Supplemental File 2:** Transformation-based dendrogram (TD) generated using the UPGMA hierarchical clustering method.

**Supplemental File 3:** Transformation-weighted characteristics dendrogram (TWCD) generated using the UPGMA hierarchical clustering method.

**Supplemental File 4:** Database of known transformations used during the transformation analysis.

**Supplemental Figure 1:** Dendrograms and sub-dendrograms with associated metrics. Panels *a*, *c*, and *e* are the visualizations of the Molecular Characteristics Dendrogram (MCD), Transformation-based Dendrogram (TD), and Transformation-Weighted Characteristics Dendrogram (TWCD), respectively. Panels b, d, and f contain sub-dendrograms of the 16 most frequently observed metabolites to demonstrate the clustering patterns of elemental composition and two metrics: average transformation count and nominal oxidation state of carbon (NOSC). Average transformation count is the average number of transformations associated with a given metabolite across all of the samples. NOSC is a metric that represents the thermodynamic availability of metabolites.

**Supplemental Document 1:** This document extensively details the materials and methods used throughout this manuscript, as well as additional discussion regarding the three metabolite dendrograms.

