## Supplemental Document 1 for "Unification of Environmental Metabolomics with Metacommunity Ecology"

**<u>Supplementary Information for:</u>**

Unification of Environmental Metabolomics with Meta-Community Ecology

### Supplemental Discussion

*Metabolite Dendrograms*

In order to investigate metabolite clustering patterns on the three dendrograms, we examined a subset of the 16 most frequently occurring metabolites within the dataset (i.e., those metabolites which appeared in the most samples) (**Supplemental Figure 1**). Furthermore, we discuss the costs and benefits of each dendrogram and their potential interpretations. All dendrograms generated during this study are provided in the supplemental material (**Supplemental File 1-3**).

<u>Molecular Characteristics Dendrogram (MCD)</u>

Following principles similar to those established in compound classification studies (Kim *et al.* 2003; Bailey *et al.* 2017; Rivas-Ubach *et al.* 2018), identified metabolites can be grouped based upon their molecular characteristics. We specifically used elemental composition (e.g., C-, H-, O-, N-, S-, P-content), double-bond equivalents (DBE), modified aromaticity index ($AI_{Mod}$), and Kendrick's defect, which all provide information regarding the composition and structure of metabolites (i.e., saturated vs. unsaturated, nitrogen content, etc.) (Hughey *et al.* 2001; Koch & Dittmar 2006; LaRowe & Van Cappellen 2011; Tfaily *et al.* 2015). These metrics were combined to estimate a pairwise Euclidean distance matrix. UPGMA (unweighted pair group method with arithmetic mean) hierarchical clustering was subsequently used to generate a dendrogram approximating compositional similarities across metabolites (**Supplemental Figure 1ab**). This dendrogram is referred to as the Molecular Characteristics Dendrogram (MCD).

The MCD appeared to provide reasonable and expected patterns of clustering. This is made clear by examining the sub-dendrogram of 16 of the most frequently observed metabolites across our sample set, within which discrete clusters formed based on metabolite characteristics. For example, metabolites containing $O_3S$ (i.e., $C_{17}H_{28}O_3S$, $C_{16}H_{26}O_3S$, $C_{18}H_{30}O_3S$) clustered together, suggesting that this feature is a dominating trait (**Supplemental Figure 1b**). In addition, more complex metabolites (i.e., those containing a mixture of O, N, S, and P, like $C_9H_3N_2O_8SP$) clustered separately from simpler CHO/CHON metabolites (**Supplemental Figure 1b**).

From this initial analysis the MCD appears to provide interpretable clustering patterns that make conceptual sense. We propose that the MCD can be used in analyses that are directly analogous to phylogenetic and functional trait analyses within community ecology. An important caveat is that while straightforward to generate and interpret, the MCD requires that metabolites have assigned chemical formulas. This requirement can lead to large portions of data being dropped. For example, within this FTICR-MS dataset, ~13% of resolved metabolites (i.e., peaks in the mass spectra) were assigned a formula. Analyses based on an MCD will, therefore, exclude a significant portion of metabolites. However, we contend that this should not preclude use of MCDs as important insights are commonly revealed through analyses of metabolite subsets that have assigned formulas (Boye *et al.* 2017; Dalcin Martins *et al.* 2017; Graham *et al.* 2017, 2018; Tolić *et al.* 2017; Stegen 2018).

<u>Transformation-based Dendrogram (TD)</u>

To provide an approach that is complementary to the MCD and that makes use of more of the resolved metabolites, we developed a dendrogram method based on inferred biochemical transformations occurring among metabolites. This is referred to as the Transformation-based Dendrogram (TD) and is enabled by the ultrahigh mass resolution of FTICR-MS, which allows all resolved metabolites with or without assigned formulas to be organized in a putative metabolic network based upon a database. This network is estimated through a series of putative biochemical transformations that are inferred through quantitative estimation of between-metabolite mass differences (Breitling *et al.* 2006; Bailey *et al.* 2017; Graham *et al.* 2017, 2018; Moritz *et al.* 2017; Stegen *et al.* 2018). For example, if the mass difference between two metabolites is 18.0343 that would indicate that there is a loss or gain of an ammonium group, while a mass difference of 103.0091 would indicate loss or gain of a cysteine. However, both the reactant and product must be present for a transformation to be detected; if one is missing, no transformation will be considered. Lists of known biochemical transformations have been previously compiled and used in analyses of ecosystem metabolomes (**Supplemental File 4**; Graham *et al.* 2018; Stegen *et al.* 2018).

To estimate the TD, we quantified the minimum number of steps a given metabolite must take to reach another metabolite in the network (e.g., metabolite 1 can become metabolite 2 via a given set of transformations). Doing so across all pairwise metabolite comparisons provides quantitative relational information among many metabolites in a given dataset (**Figure 1**). In turn, the TD is estimated from the pairwise metabolite network distances and is therefore based on information of how each metabolite could give rise to another (**Supplemental Figure 1cd**).

The topology of the TD can diverge from the MCD because it is based on putative biochemical transformations that link metabolites, rather than metabolite molecular characteristics (**Supplemental Figure 1d**). For example, in the MCD there were high O-content (i.e., $C_9H_2O_{10}$, $C_{10}H_2O_{11}$) and low O-content (i.e., $C_{17}H_{26}O_5$, $C_{18}H_{34}O_3$, etc.) clusters that diverged from each other, but were relatively near each other in the TD. This is likely because there are a small number of transformations (i.e., network steps) separating metabolites with high O-content from the low O-content metabolites. In addition, other metabolite-specific variables such as average transformation count, which measures the average number of transformations associated with a given peak, are more strongly associated with the topology of the TD than the MCD (**Supplemental Figure 1d**). This points to structure within the network whereby highly connected metabolites are linked to other highly connected metabolites through a small number of biochemical transformations. This type of non-random structure within the broader transformation network is captured in the TD and therefore informs downstream evaluation of processes governing the spatiotemporal organization of metabolite assemblages.

Importantly, the TD can be generated without formula assignments. This allows a greater number of metabolites to be used in and inform other analyses, which is an advantage over the loss of data that occurs with the MCD. While this increase in peak representation can be beneficial, it does come at the cost of not considering metabolite characteristics that provide biogeochemically-relevant stoichiometric information. Therefore, the TD and MCD should be viewed as complementary. There are likely scenarios in which analyses based on TD and MCD would be used together to deepen insights.

Transformation-Weighted Characteristics Dendrogram (TWCD)
While the MCD and TD each have unique strengths, there is an opportunity to combine these approaches to obtain more information than is available with either on its own. To do so, among metabolite Euclidean distances based on molecular characteristics were weighted by among-metabolite distances derived from the biochemical transformation network. In effect, the combination of these two matrices results from the transformation-based matrix (standardized from 0 to 1) decreasing the characteristics-based distance values. The resulting dendrogram is referred to as the Transformation-Weighted Characteristics Dendrogram (TWCD) (**Supplemental Figure 1e**). Within the TWCD, metabolites that are distinct based on their characteristics but that are close in the transformation network should cluster more closely together than in the MCD but further apart than in the TD. Such changes can be observed in the TWCD sub-dendrogram in which the $O_3S$ cluster is closer to the low O-content cluster than it was in the MCD (**Supplemental Figure 1f**), while also being further from the single $OS_2$ compound than it was in the TD. As another example, the low O-content cluster in the TWCD (e.g., metabolites with $O \leq 3$) is further from the other CHO-metabolites than it was in either the MCD or TD. This suggests that the molecular characteristics of low O-content metabolites are more different from CHO-metabolites than would be expected given the connecting biochemical transformations between them.

We suggest that by merging molecular characteristics with biochemical transformations, TWCDs carry more information related to functionally relevant attributes of metabolites, relative to MCDs or TDs. Despite merging information from MCDs and TDs, however, analyses performed with TWCDs will not be as sensitive to differences in either molecular characteristics or the biochemical transformation network. This again indicates an opportunity to draw out deeper insights by combining patterns derived from multiple dendrograms. We therefore suggest that while there are other approaches to estimating dendrograms from metabolite data, the MCD, TD, and TWCD provide a complementary set of analysis tools that are useful for studying the spatiotemporal organization of metabolite assemblages.

**Supplemental Materials and Methods**
*Sample Collection.* River and pore water samples were collected from the Columbia River in Washington State along a ~1 km transect along the shoreline. At each location, one replicate of river water was collected, and 3 pore water samples were collected and filtered using a 0.2 µm Sterivex filters (MilliporeSigma, MA, USA). Pore water replicates were collected from 30cm depth within a 1m$^2$ area using 0.25-inch diameter sampling tubes (MHE Products, MI, USA). Filters were stored at -80 C until DNA could be extracted while water samples were stored at -20C until they could be used for further analysis.

*DNA Extraction, Sequencing, and Processing.* DNA was extracted from Sterviex filters using a Powersoil DNA isolation kit (Mo Bio Laboratories, Inc., Carlsbad, CA). In order to generate 16S rRNA gene data, the V4 region of 16S rRNA genes was amplified and sequenced using the universal bacterial/archaeal primer set 515F/806R on an Illumina MiSeq instrument at Argonne National Laboratory according to the Earth Microbiome Project standard protocol (Caporaso *et al.* 2012). Resulting 16S rRNA amplicon sequences were analyzed using the open access 'hundo' pipeline (Brown *et al.* 2018). Adapters were trimmed, low quality reads (i.e., length < 100 bp, quality score > 10), and contaminant sequences were filtered using BBDuk2 from the BBTools package (Bushnell 2018). Reads passing the quality filter were then clustered at 97% into *de novo* operational taxonomic units (OTUs) using VSEARCH with a minimum merge length of 150 bp and a minimum sequence abundance of 2 (Rognes *et al.* 2016). Simultaneously, chimeric sequences were removed through *de novo* prediction and reference-based identification. Following clustering, BLAST was used to align sequences to the SILVA nr SSU reference database (Camacho *et al.* 2009; Quast *et al.* 2013) and taxonomy was assigned based upon the CREST lowest common ancestor classifier (Lanzén *et al.* 2012). Sequences were aligned using Clustal Omega (Sievers *et al.* 2011) and a phylogenetic tree was generated using FastTree 2.0 (Price *et al.* 2010).

*Fourier Transform Ion Cyclotron Resonance Mass Spectrometry Sample Preparation, Data Collection, and Data Preprocessing.* Fourier Transform Ion Cyclotron Resonance Mass Spectrometry (FTICR-MS) was used for the ultrahigh resolution characterization of dissolved organic matter (DOM) within each sample. Filtered river and pore water samples were acidified to pH 2 with 85% phosphoric acid and extracted with PPL cartridges (Bond Elut), following Dittmar et al. (e.g., solid-phase extraction) (Dittmar *et al.* 2008). High-resolution mass spectra of the DOM were collected using a 12 Tesla (12T) Bruker SolariX Fourier transform ion cyclotron resonance mass spectrometer (Bruker, SolariX, Billerica, MA) located at the Environmental Molecular Sciences Laboratory in Richland, WA. Samples were directly injected into the instrument using a custom automated direct infusion cart that performed two offline blanks between each sample. The FTICR-MS was outfitted with a standard electrospray ionization (ESI) source, and data was acquired in negative mode with the needle voltage set to +4.4kV, resolution was 220K at 481.185 m/z. One hundred forty-four scans were co-added for each sample and internally calibrated using organic matter homologous series separated by 14 Da (–CH2 groups). The mass measurement accuracy was typically within 1 ppm for singly charged ions across a broad m/z range (100 m/z - 900 m/z). The FTMS peak picker module in the BrukerDaltonik Data Analysis software (version 4.2) was used to convert raw spectra to a list of m/z values with a signal-to-noise ratio (S/N) threshold set to 7 and absolute intensity threshold to the default value of 100. Formularity (Tolić *et al.* 2017), an in-house software, was used to assign chemical formulae following the Compound Identification Algorithm (Kujawinski & Behn 2006) and to align peaks with a 0.5 ppm threshold. Chemical formulae were assigned based on the following criteria: S/N >7, and mass measurement error < 0.5 ppm, taking into consideration the presence of C, H, O, N, S and P and excluding other elements.

The R package ftmsRanalysis (Bramer & White 2019) was used to remove peaks that either were outside the desired m/z range (200 m/z – 900 m/z) or had an isotopic signature, calculate derived statistics (Kendrick defect, double-bond equivalent, aromaticity index, nominal oxidation state of carbon, standard Gibb's Free Energy of carbon oxidation), and organize the data (Hughey *et al.* 2001; Koch & Dittmar 2006; LaRowe & Van Cappellen 2011; Tfaily *et al.* 2015). Given that charge competition renders peak intensities less informative across systems (Tfaily *et al.* 2017), all analyses were conducted using binary presence/absence values rather than peak intensities with the absence of a peak defined as being below the limit of detection.

*Metabolite Dendrogram Estimation.* Three different metabolite dendrograms were generated: the molecular characteristics dendrogram (MCD), transformation-based dendrograms (TD), and transformation-weighted characteristics dendrogram (TWCD). Using the derived statistics calculated above (e.g., elemental composition, double-bond equivalents, modified aromaticity index, and Kendrick's defect), we can compare the potential molecular similarities between identified chemical formulae as in compound classification studies (Kim *et al.* 2003; Bailey *et al.* 2017; Rivas-Ubach *et al.* 2018). Multivariate similarities were evaluated by measuring the Euclidean distance between chemical formulae (*vegdist*, vegan package v2.5-6) (Oksanen *et al.* 2019). These Euclidean distances were then used to perform a UPGMA hierarchical cluster analysis (*hclust*, 'average' method, stats package).

Unlike the MCD, the TD estimates molecular similarity by inferring potential biochemical transformations based upon ultrahigh mass resolution differences between identified metabolites (Breitling *et al.* 2006; Graham *et al.* 2017, 2018; Moritz *et al.* 2017; Sengupta *et al.* 2019). For example, if the mass difference between two metabolites was 18.0343, that would putatively indicate a loss or gain of an ammonium group, while a mass difference of 103.0092 would putatively indicate loss or gain of a cysteine. This calculation is enabled by the ultrahigh mass resolution of FTICR-MS data; given this resolution, using a transformation database (**Supplemental File 4**), we considered any between-metabolite mass difference within 1 ppm of the expected mass of a transformation to be a match. Using these pairwise mass differences and transformation associations, we generated a transformation network in which nodes represent individual metabolites and edges are identified transformations. Relationships between metabolites were determined by selecting the largest cluster of interconnected nodes (discarding any node outside this cluster) and measuring the stepwise distance between each pair of metabolites (i.e., the minimum number of transformations required to connect one metabolite to another metabolite within the largest cluster of the biochemical transformation network). These pairwise distances were then standardized between 0 and 1. A UPGMA hierarchical cluster analysis (*hclust*, 'average' method, stats package) was then used to convert these distances into a dendrogram.

The TWCD is a composite dendrogram requiring partial creation of both the MCD and TD. First, a Euclidean molecular characteristics distance matrix must be created based upon the elemental composition and derived statistics (i.e., double-bond equivalents, etc.). Next, the standardized stepwise transformation distance matrix (i.e., values between 0 and 1) must be generated based upon the transformation analysis described above. Using simple matrix multiplication, the molecular characteristics matrix is combined with the standardized transformation matrix. In effect, this results in a matrix where realized molecular characteristic differences are down-weighted. The TWCD is the generated by performing a UPGMA hierarchical clustering analysis on this transformation-weighted, molecular characteristics distance matrix.

α- *and* β-*diversity analyses.* Taxonomic richness was calculated by counting the total number of metabolites in each sample. Numerous dendrogram-based (i.e., phylogenetic) α-diversity measurements were also utilized within this study and are more completely detailed within Tucker *et al.* (2016). In brief, three dendrogram-based α-diversity metrics were utilized: Faith's Phylogenetic Diversity (PD; here, referred to as Dendrogram Diversity or DD) (Faith 1992), mean pairwise diversity (MPD) (Webb *et al.* 2002), and mean nearest taxon distance (MNTD) (Webb *et al.* 2002). Faith's PD (or DD) is a richness measurement that sums the total length of dendrogram branches that connect taxon:

$$PD \text{ or } DD = \sum_{b \in B} L_b$$

where $b$ is a given branch with a larger set of branches $B$ and $L_b$ is the length of branch b. MPD is a metric which determines the average dendrogram distance between taxon:

$$MPD = \frac{\sum_{ij} d_{ij}}{S(S-1)}$$

where $d_{ij}$ is the distance between taxon $i$ and taxon $j$ and $S$ is the total number of species within the community. Lastly, MNTD is a metric which determines the average dendrogram distance between nearest neighbors (i.e., the next closest taxon on the dendrogram):

$$MNTD = \frac{1}{S}\sum_{i} d_{i\,min}$$

where $d_{i\,min}$ is the shortest distance from taxon $i$ to all other taxon. Faith's PD was calculated using the *pd* function in the picante R package (v1.8) (Kembel *et al.* 2010) while MNTD and MPD were measured using the *generic.metrics* function in the pez R package (v1.2-0) (Pearse *et al.* 2015). These metrics were calculated using each of the three dendrograms.

Taxonomic β-diversity was visualized by generating a Jaccard dissimilarity-based non-metric multidimensional scaling (NMDS) ordination (*metaMDS*, vegan package v2.5-6) (Oksanen *et al.* 2019). Using each of the dendrograms, dendrogram-based (i.e., phylogenetic) beta-diversity was calculated using the unweighted UniFrac metric (*GUniFrac*, GUniFrac package v1.1) (Chen 2012). UniFrac results were visualized using a principal coordinate analysis (PCoA; *pcoa*, ape package v5.3) (Paradis & Schliep 2019).

*α- and β-diversity ecological null modeling.* Both α- and β-diversity based null modeling was performed throughout this study. In order to assess whether α-diversity was more or less structured than would be expected by random chance, both net relatedness index (NRI) and nearest taxon index (NTI) were calculated (Webb *et al.* 2002; Fine & Kembel 2011; Kraft *et al.* 2011). NRI is the null model variant of MPD and measures the degree of dispersal across the entire dendrogram, while NTI is the null model variant of MNTD and identifies tip-level clustering. For each of these null model calculations, 999 randomized metabolite assemblages were generated through tip shuffling. The finalized index was then calculated:

$$NRI = -1\left(\frac{MPD_{obs} - \overline{MPD_{null}}}{MPD_{sd}}\right)$$

$$NTI = -1\left(\frac{MNTD_{obs} - \overline{MNTD_{null}}}{MNTD_{sd}}\right)$$

where $MPD_{obs}$ and $MNTD_{obs}$ are the observed α-diversity values, $\overline{MPD_{null}}$ and $\overline{MNTD_{null}}$ are the average α-diversity values derived from the null assemblages, and $MPD_{sd}$ and $MNTD_{sd}$ are the standard deviations of α-diversity values from the null assemblages. Both NTI and NRI can be interpreted similarly; positive values suggest phylogenetic clustering while negative values indicate phylogenetic overdispersion (Webb *et al.* 2002).

β-diversity null modeling was performed to investigate whether metabolite assemblages were significantly more or less similar than would be expected by random chance alone and to assess whether assemblages were deterministically or stochastically assembled. To explore these ecological assembly processes, we first calculated the dendrogram-based β-nearest taxon index (βNTI) for each possible pairwise comparison according to 10. First, β-mean nearest taxon index (βMNTD) for the observed metabolite assemblages must be calculated in order to estimate dendrogram-based turnover:

$$\beta\text{MTND} = \frac{\sum_{i_k=1}^{n_k} f_{i_k} \min\left(d_{i_k j_m}\right) + \sum_{i_m=1}^{n_m} f_{i_m} \min\left(d_{i_m j_k}\right)}{2}$$

where $f_{i_k}$ is the relative abundance of metabolite *i* in community *k*, $n_k$ is the number of metabolites in community k, and min $(d_{i_k j_m})$ is the minimum dendrogram distance between metabolite *i* in community $k$ and metabolite $j$ in community $m$. This metric was calculated using the *comdistnt* function (abundance.weighted = FALSE) in the picante R package (v1.8) (Kembel *et al.* 2010). Similar to the NRI and NTI calculations, 999 randomized communities were generated by shuffling the tips of the dendrogram. βMNTD was then determined for each of these null communities and βNTI was calculated:

$$\beta\text{NTI} = -1\left(\frac{\beta\text{MTND}_{obs} - \overline{\beta\text{MTND}_{null}}}{\beta\text{MTND}_{sd}}\right)$$

where $\beta\text{MTND}_{obs}$ is observed βMNTD for the observed assemblages, $\overline{\beta\text{MTND}_{null}}$ is the average βMNTD for the null communities, and $\beta\text{MTND}_{sd}$ is the standard deviation of $\beta\text{MTND}_{null}$ values. In order to compare the different dendrogram estimation methods, we independently calculated βNTI values for each of the three metabolite dendrograms (e.g., MCD, TD, and TWCD). Microbial βNTI values were generated using the 16S rRNA gene amplicon phylogenetic tree. We also calculated Resulting βNTI values help examine phylogenetic turnover across samples, providing insight into the ongoing deterministic and stochastic ecological assembly processes occurring within the system. If a |βNTI| value is greater than 2, deterministic processes explain the observed assemblage differences; if a |βNTI| value is less than 2, stochastic processes are responsible for assemblage differences. Stochastic processes are those which are more random in nature and typically arise from dispersal-based events. Deterministic processes are those which are driven by environmental filtering, pushing assemblages to either be more or less similar than expected by random chance Deterministic processes could be further broken down into variable selection if βNTI is greater than 2, and homogenous selection if βNTI is less than -2. Variable selection occurs when the environment drives assemblages to be significantly divergent, as observed when distinct geochemistry supports different microbial communities (Danczak *et al.* 2018). In contrast, homogeneous selection occurs when some common pressure push communities toward a similar configuration as has been observed in microbial communities experiencing salt stress in a soil succession system (Dini-Andreote *et al.* 2015). Correlations between microbial and metabolite βNTI values were performed by averaging the βNTI values for a given assemblage and relating them within a sample.

In addition to the dendrogram-based βNTI, we distinguished stochastic processes by using the identity-based Raup-Crick (RC) (Stegen *et al.* 2013, 2015). Using 9,999 iterations per pairwise comparison, null communities were probabilistically generated based upon observed metabolite assemblages and presence/absence-based Bray-Curis (i.e., Sørensen) dissimilarities were calculated. The null distribution of these dissimilarity values was then compared to the observed Bray-Curtis value in order to measure the deviation from the null expectation. These deviations were then standardized to vary between -1 and 1, resulting in the finalized RC metric. If a |RC| value is greater than 0.95, the turnover between the compared assemblages was the result of either dispersal limitation or homogenizing dispersal. Dispersal limitation (RC > 0.95) occurs when assemblages are unable to mix resulting in significant ecological drift. Conversely, homogeneous dispersal (RC < -0.95) occurs when environments drive substantial mixing resulting in assemblages that are more similar than random chance alone. However, if a |RC| value is less than 0.95, the assemblages were as different as would be expected by random chance because no single process is able to dominate (i.e., weak selection and weak dispersal). Under these 'undominated' circumstances, no single assembly process is capable of dominating. Significant differences in distributions of both βNTI and RC values across surface and pore water classifications were identified using Mann Whitney U tests (*wilcox.test*, stats package).

*Metabolite Activity Comparisons.* In order to identify putatively more active metabolites, we leveraged potential biochemical transformation information collected during the generation of the transformation-based dendrogram (TD). Specifically, we counted the number of transformations associated with each individual metabolite and arbitrarily decided that metabolites involved in more than 40 transformations were considered 'active.' Conversely, metabolites involved in no transformations were considered 'inactive.' Given that our transformation database is not exhaustive, we caution that these classifications are not generalizable but serve as a sufficient exercise. Using these two metabolite sub-assemblages (e.g., active and inactive), we calculated βNTI for all three of the metabolite dendrograms by pruning the dendrograms to match the data.

*Plot Generation.* All box plots and pie charts were generated using ggplot2 (Wickham 2016). Dendrograms and their associated bar charts were visualized using ggtree (Yu *et al.* 2017).

All scripts and raw data used in the realization of this manuscript are hosted on GitHub at https://github.com/danczakre/Meta-Metabolome_Ecology.

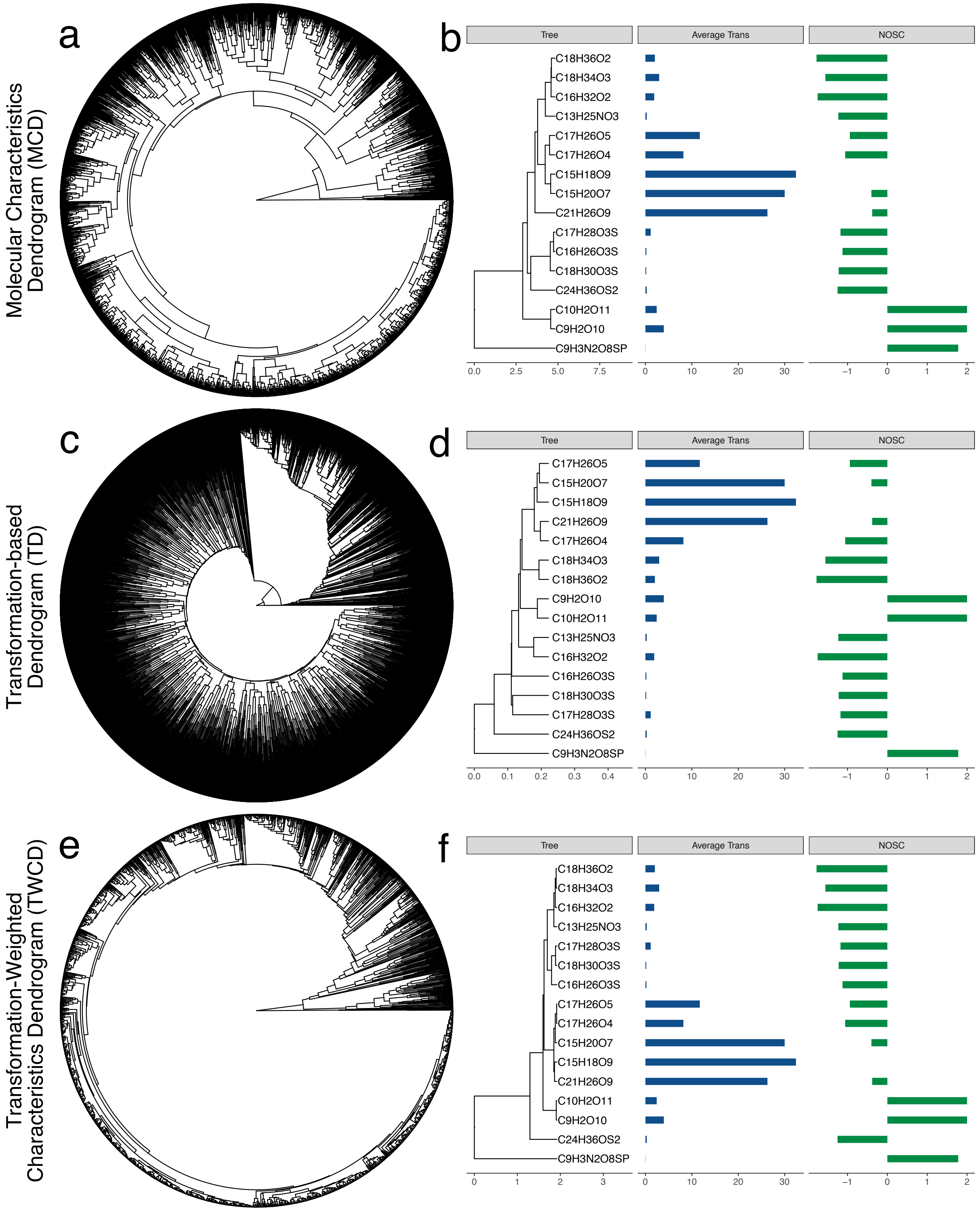


**Supplemental Figure 1:** Dendrograms and sub-dendrograms with associated metrics. Panels *a*, *c*, and *e* are the visualizations of the Molecular Characteristics Dendrogram (MCD), Transformation-based Dendrogram (TD), and Transformation-Weighted Characteristics Dendrogram (TWCD), respectively. Panels b, d, and f contain sub-dendrograms of the 16 most frequently observed metabolites to demonstrate the clustering patterns of elemental composition and two metrics: average transformation count and nominal oxidation state of carbon (NOSC). Average transformation count is the average number of transformations associated with a given metabolite across all of the samples. NOSC is a metric that represents the thermodynamic availability of metabolites.
